# Odour conditioning of positive affective states: Rats can learn to associate an odour with being tickled

**DOI:** 10.1101/549436

**Authors:** Vincent Bombail, Nathalie Jerôme, Ho Lam, Sacha Muszlak, Simone L Meddle, Alistair B Lawrence, Birte L Nielsen

## Abstract

Most associative learning tests in rodents use negative stimuli, such as an electric shock. We investigated if young rats can learn to associate the presence of an odour with the experience of being tickled (i.e. using an experimenter’s hand to mimic rough-and-tumble play), shown to elicit 50 kHz ultrasonic vocalisations (USVs), which are indicative of positive affect. Male, pair-housed Wistar rats (N=24) were all exposed to two neutral odours (A and B) presented in a perforated container on alternate days in a test arena. Following 60s of exposure, the rats were either tickled on days when odour A (n=8) or odour B (n=8) was present, or never tickled (n=8). When tickled, rats produced significantly more 50 kHz USVs compared to the days when not being tickled, and compared to control rats. The level of anticipatory 50 kHz USVs in the 60s prior to tickling did not differ significantly between the tickled and control rats. Following the odour conditioning, rats were exposed successively in the same arena to three odours: an unknown neutral odour, extract of fox faeces, and either odours A or B. Compared to controls, 50 kHz USVs of tickled rats increased when exposed to the odour they had previously experienced when tickled, indicating that these rats had learned to associate the odour with the positive experience of being tickled. In a test with free access for 5 min to both arms of a T-maze, each containing one of the odours, rats tickled with odour A spent more time in the arm with this odour. This work is the first to test in a fully balanced design whether odours can be conditioned to tickling, and indicates that positive odour conditioning has potential to be used as an alternative to negative conditioning tests.

## Introduction

Aversive conditioning, where a previously neutral stimulus or place is associated with an aversive stimulus, can be used to study memory and other brain functions in laboratory rodents (e.g. Ellis and Kesner, 1983). This paradigm is used in studies of learning, as odours can be associated with aversive states such as fear (Kroon and Carobrez, 2009; Moriceau et al., 2006) and malaise (Chapuis et al., 2007; Raineki et al., 2009). The animals usually learn this association very quickly, making it a time-saving and efficient research method, which may be why only few attempts have been made to develop tests for this purpose using positive experiences. In studies where positive conditioning of odours has been applied, they involved pairing with psychostimulant drugs (Revillo et al., 2012; Caffrey and Febo, 2014; Lowen et al., 2015), alcohol (Deehan et al., 2012) or a food source (Shide and Blass, 1991; Sullivan et al., 2015; Torquet et al., 2014). However, using feed as the unconditioned stimulus is not always feasible in practice, and is likely to be associated with an increasing level of satiety.

We were therefore interested in finding an appropriate positive conditioning stimulus for use in an associative learning test for rats. This would ideally consist of a stimulus that was easy to use and which gave rise to the animal experiencing positive welfare (i.e. a positive affective state and not just absence of negative welfare; Lawrence et al., 2017). Despite the increasing interest in positive welfare indicators, the vast majority of animal welfare research has been and continues to be focused on more negative aspects (e.g. Boissy et al., 2007). One result of this is that there are few well-validated models of positive welfare in animals. One of the best candidates is the rat tickling model that was developed to mimic the effects of social play, a behaviour frequently displayed by young rats (see LaFollette et al. (2017) for a recent review). In this model, the human hand is used to mimic the tactile stimulation experienced during social play in rats. The model has been validated partly through the measurement of ultrasonic vocalisations (USVs) that rats produce under different emotional states. During tickling, rats produce many more frequency modulated USVs in the range of 33-100 kHz (henceforth referred to as 50 kHz USVs); these are sometimes referred to as ‘laughter’ (Panksepp and Burgdorf, 2010), and have been shown to indicate a positive emotional state (Knutson et al., 2002). Tickled rats show shorter latencies to approach the human hand than controls, and express so-called optimistic biases when appraising environmental cues (Rygula et al., 2012). In addition, a number of pharmacological manipulations of rats using various psychotropes supports the notion that 50 kHz USVs are produced upon activation of the brain’s reward pathways (Popik et al., 2014; Avvisati et al., 2016). Data thus support the interpretation that expressions of 50 kHz USVs indicate that tickling is a positive experience for the rat. However, USVs in the range of 22 kHz are emitted by rats under aversive situations (Tonoue et al., 1986; Blanchard et al., 1991; Choi and Brown, 2003; Burgdorf et al., 2018). USVs have therefore been suggested to be a useful tool for inferring affective states of the rats (Burgdorf and Panksepp, 2006; Burgdorf et al., 2011; Wöhr and Schwarting, 2013; Barker, 2018).

In this paper, we present the results of an experiment which was designed to condition rats to associate the presence of an odour with the positive experience of tickling. We hypothesised that if rats learned to make the odour-tickling association, they would i) emit anticipatory USVs when exposed to the odour prior to being tickled, *ii)* emit more 50 kHz USVs than control rats when exposed to the conditioned odour following exposure to an aversive odour, and *iii)* would spend more time in the arm of a T-maze containing their tickling odour.

## Materials and Methods

Male Wistar rats (n=24) were used as subjects for odorant conditioning and subsequent behavioural testing. The rats were housed in pairs at 4 weeks of age in standard laboratory rodent cages (42.5 cm × 26.6 cm × 18.5 cm made from transparent polycarbonate; Techniplast 1291H) on a 4-tier rack. The lighting schedule of the room was inverse 12D:12L, with lights coming on at 19:00 hours. The cages had a metal grid lid with a dentation in which commercial rat pellets (Diet M25, Special Diet Services, Witham, Essex, United Kingdom) were placed for ad libitum access. Water was supplied via a drinking bottle with a metal spout, inverted and placed alongside the feed. The floor of the cage was covered by 2 cm of sawdust litter changed weekly, and wooden chew sticks (12 cm long) were supplied as enrichment. Individual rats were identified by marker pen lines on the tail. The rats were weighed once a week and, if needed, their tails were remarked.

The rats were handled daily by the same person, and all handling, conditioning and testing took place during the dark period. Over the course of 5 days, the rats were gradually habituated to being put into a transport box (identical to the home cage, but with no water and feed available), and transported within an opaque black sack to the conditioning room, which was illuminated by red incandescent bulbs. The rats were also habituated to the conditioning arena – initially in pairs and subsequently individually. The arena consisted of a Plexiglas tank (L × W × H: 66 cm × 41 cm × 41 cm), bedded with sawdust. At one end of the arena, a thin metal plate was fixed centrally at the bottom of the wall. During habituation, an empty stainless steel container (diameter 9.5 cm; height 3.7 cm; Grundtal IKEA) with a magnetic base and a screw-top lid with a perforated plastic inset was affixed vertically to the metal plate (Figure 1a).

**Figure 1.**
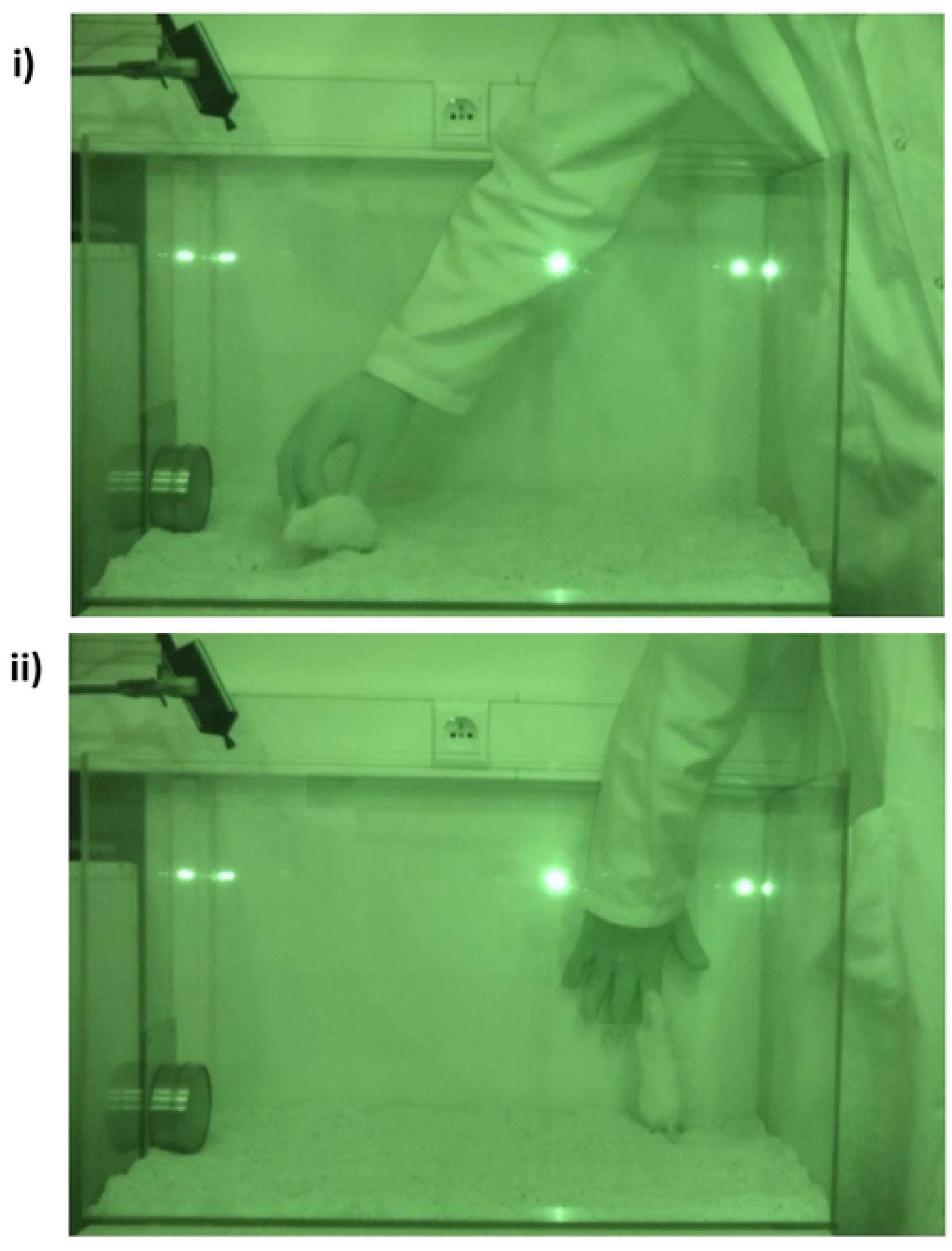
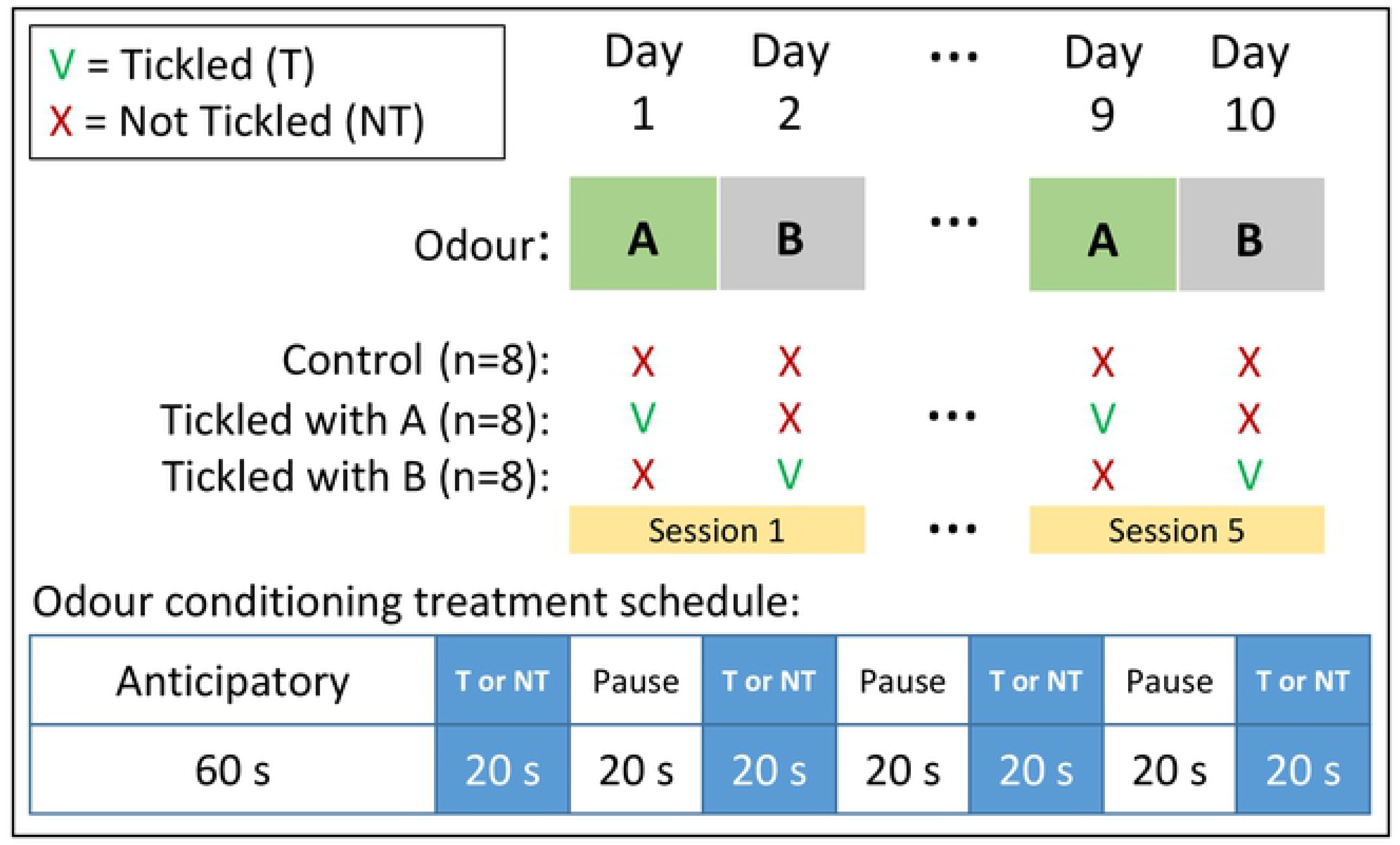
**a)** Screen-shots from the video recording of an odour conditioning session, showing **i)** the rat being tickled, and **ii)** the position of the hand flat against the arena wall during the pauses between tickling periods. The container with the odour source can be seen in the left of the pictures. The pictures have a green hue as the procedure was carried out under red lighting; **b)** Diagram of the odour conditioning schedule over days and within treatment. Control rats were never tickled and A-tickled rats were tickled when odour A was present in the container, and B-tickled rats were tickled when odour B was present in the container in the test arena. When odour A was in the container, the B-tickled rats were treated as the Control rats, as was the A-tickled rats when odour B was in the container.

In order to account for potential odour dependent effects, two different odours were used as the conditioning odours: **Odour A** (a 10% dilution of D-limonene; CAS no. 5989-27-5) and **Odour B** (a 5% dilution of 1-hexanol; CAS no. 111-27-3). Both were diluted in mineral oil (CAS no. 8042-47-5) at different concentrations to obtain similar levels of odour strength as assessed by the researchers, and 2 ml samples were transferred to a cotton pad in the container prior to the conditioning. All compounds were purchased from Sigma Aldrich (Saint-Quentin, Fallavier, France).

Each rat was allocated to one of three conditioning treatments, and all rats were placed in the arena on every conditioning day, where its conditioning treatment was applied: **Tickled A** rats were tickled when odour A was present in the container in the test arena; **Tickled B** rats were tickled when odour B was present in the container in the test arena; and **Control** rats were never tickled but still presented with either of the odours in the arena on alternate days (Figure 1b). When odour A was in the container, the Tickled B rats were treated as the Control rats, as were the A-tickled rats when odour B was in the container. Each pair of rats within a cage was randomly allocated to a treatment, whilst ensuring that rats housed on each tier of the rack received all three treatments. The order in which the rats were treated changed from one day to the next, starting with rats 1, 19, 13, and 7 on different days, respectively.

Conditioning began when the rats were 6 weeks of age. Only one of the two odours were used on each conditioning day to minimise the risk of cross-contamination. All conditioning sessions where video recorded (Sony 12.0 mega pixels HDR-XR-500 Handycam) and the USVs registered using a free-ware sound-recording programme (Audacity 2.1.3; www.audacityteam.org) via a USV sensitive microphone (M500-384, Pettersson Elektronik, Sweden) attached to the arena at an angle of 45° above the container (Figure 1a). A session began by moving the rat from its home cage in the transport box to the conditioning room and placing the rat in the arena 10 cm from and facing the container containing one of the two odours, A or B. The rat was left there for 1 min, and the rat was subsequently either **Tickled** (Tickled A rats on days when odour A was present, Tickled B rats on days when odour B was present) or **Not Tickled** (Control rats on all days, Tickled A rats when odour B was present, and Tickled B rats when odours A was present; Figure 1b) according to the following procedures:

**Tickled** consisted of the handler using one hand wearing a knitted glove to touch, tickle, and play with the rat for 20 second periods interspersed with 20 second pauses. During the active periods, the handler mimicked the rough-and-tumble play seen in adolescent rats, with the hand tickling, chasing and pinning the rat, depending on its response (Figure 1a.i). After 20 seconds the hand was placed flat on the inside of the wall of the arena (Figure 1a.ii). It rested here for 20 seconds after which another tickling period was carried out. This was repeated for a total duration of 140 s, allowing 4 periods of tickling interspersed with 3 periods of pauses. At the end of the tickling, the rat was moved in the transport box to a holding cage in a room separate from their home cage. The holding cage was identical to the home cage of the rat, placed in a similar 4-tier rack in the same position, and one holding cage was used for each pair of rats. The tickled rats were left in the holding cage for 3-7 hours (depending on the test order of the day) to prevent emotional contagion by USVs of the yet un-tested rats in the home cages.

**Not Tickled** consisted of the handler placing the hand wearing a knitted glove flat on the inside of the wall of the arena. It rested here for 20 seconds after which the hand was moved to the adjacent wall for 20 seconds. This was repeated for a total duration of 140 s, allowing 4 and 3 periods, respectively, with the hand resting on each wall, the latter being identical to the pauses when the rats were being Tickled. When the Not Tickled procedure finished, the rat was moved in the transport box back to its home cage.

LaFollette et al. (2018) found that three tickling sessions sufficed to bring about a higher rate of 50 kHz USVs in tickled rats, and in a pilot study using only one odour, we found a significant difference in 50 kHz USVs emitted between tickled and control rats after four tickling days (Lam, 2017). Nevertheless, as two odours were used alternately in the present experiment, we chose to carry out a total of ten conditioning days, alternating between odours A and B (Figure 1b). This resulted in all tickled rats being exposed to the Tickled procedure five times and the Not Tickled procedure five times, and all the rats were exposed to both odours for the same amount of time over the course of the ten days, including the Control rats.

For subsequent data analyses, the 50 kHz USVs for each rat were counted for the first (days 1 and 2) and the fifth (days 9 and 10) conditioning sessions, using the method described by Brenes and Schwarting (2014). These counts were divided into 50 kHz USVs emitted during the four tickling periods (4 × 20s) and the three pauses (3 × 20s), as well as the 1-min habituation period prior to tickling to detect any differences in potential anticipatory USVs. This grouping of 50 kHz USVs was also done when the Not Tickled procedure was applied (Figure 1b). All counts were converted into USVs/min. Occurrences of 22 kHz USVs were rare, and not included in the data.

The behaviour of the rats was logged from the video recordings. This was done for the three 20s pauses only, as the hand did not move and the behaviour of the rats at this time was therefore comparable within and among all rats and across treatments and procedures. The recorded behaviour consisted of hand seeking behaviour (the rat rears to sniff the motionless hand, see Figure 1a.ii, or is facing and focusing on the hand), play jumping (jumping while running), sniffing the air, exploring the odour container, freezing (immobility, often sudden, with ears raised and eyes open) and other behaviour (locomotion, digging the litter, and self-grooming).

### Behavioural tests of conditioning

On the two days following the last conditioning session, two behavioural tests were carried out to investigate the effects of the conditioning:

#### T-maze test

Without prior habituation to the T-maze, the rat was placed at one end of a large T-maze arena, which consisted of a rectangular open space (77 cm × 51 cm) with two accessible arms (WxL: 19 cm × 25 cm) extending from each side at one end of the rectangle, forming a broad T-shape. A ventilator fitted centrally at the other end of the rectangle extracted air from the T-maze, ensuring a simultaneous airflow from both arms. No litter was used, and a perforated metal tea-ball was placed in a pre-drilled hole at the end wall of each arm of the maze. The two tea-balls each contained a cotton pad imbibed with 2 ml of either odour A or B, with one odour in each arm, alternating between arms in a balanced way for each rat being tested. The test was video recorded from above. The rat was left in the arena for 3 min, and was free to explore both arms and the central arena.

#### Triple Odour test

The rat was placed in the same arena as used for the conditioning. The tests consisted of 30-sec periods with no odour container present in the arena, interspersed with three 1-min periods, where a container was positioned containing the following odours in said order: *1)* a neutral odour unknown to the rat (**Novel odour;** a 5 % suspension of p-anisaldehyde, CAS no. 123-11-5, in mineral oil), which was assumed to have no aversive or attractive properties for the rats; *2)* an extract in mineral oil of fox faeces (**Fox odour;** faecal pellets originating from several male foxes and soaked in mineral oil for 24h at 70°C, with extract diluted 1:6); this odour was expected to induce a level of fear in the rats, and *3)* the **Tickling odour** with which the Tickled rats had been conditioned, and with half of the Control rats being exposed to odour A and the other half to odour B. The order of the three odours were chosen so as to measure the response of the rats to first an unknown, but neutral odour, then an unknown but fear-inducing odour, followed by the known conditioning odour. We hypothesised that if the Tickled rats had learned to associate their tickling odour with a positive experience, more 50 kHz USVs would be emitted by the Tickled rats compared to the Control group when exposed to the conditioning odour, the latter having been exposed to the odour for the same amount of time during conditioning but without being tickled. The tests were video recorded and the amount of freezing displayed by the rats during exposure to the three different odours was scored. The USVs were registered in the same way as for the conditioning sessions.

#### Statistical data analysis

Data were analysed in MiniTab (ver. 17.1) using General Linear Models followed by post-hoc Tukey comparisons of significant effects. For the USVs emitted during conditioning, data from the four tickling periods were analysed fitting odour, treatment, and session no. with interaction. When relevant, Pearson’s correlations were calculated. Anticipatory USVs were analysed for session 5 only, fitting odour, treatment and their interaction. Behaviour during pauses, expressed as percentage of time spent on each behaviour, was analysed by the non-parametric Kruskal-Wallis test, as data were not normally distributed.

## Results

All tickled rats emitted significantly more 50 kHz USVs during the sessions with the Tickled procedure than during the sessions when Not Tickled (F_2,89_=31.3; P<0.001), with the latter not differing in magnitude from that of the never tickled Control rats (Figure 2a). This was evident already during the very first tickling session, but with significantly more 50 kHz USVs emitted during the 5^th^ compared to the 1^st^ tickling session (233 vs 83 (±8.4) USVs/min; P < 0.001), and these were significantly correlated (Pearson’s r=0.53; P = 0.036), indicating that response level of 50 kHz USVs to tickling is a characteristic of the individual rat. No differences were found between rats tickled with Odours A and B, respectively. On days when the tickled rats were Not Tickled (i.e. A-tickled rats when odour B was present and vice versa), the 50 kHz USVs emitted per minute by the 5^th^ tickling session did not differ significantly from the level observed during tickling in the 1^st^ session, indicating a degree of place association had developed (Figure 2a).

**Figure 2a.**
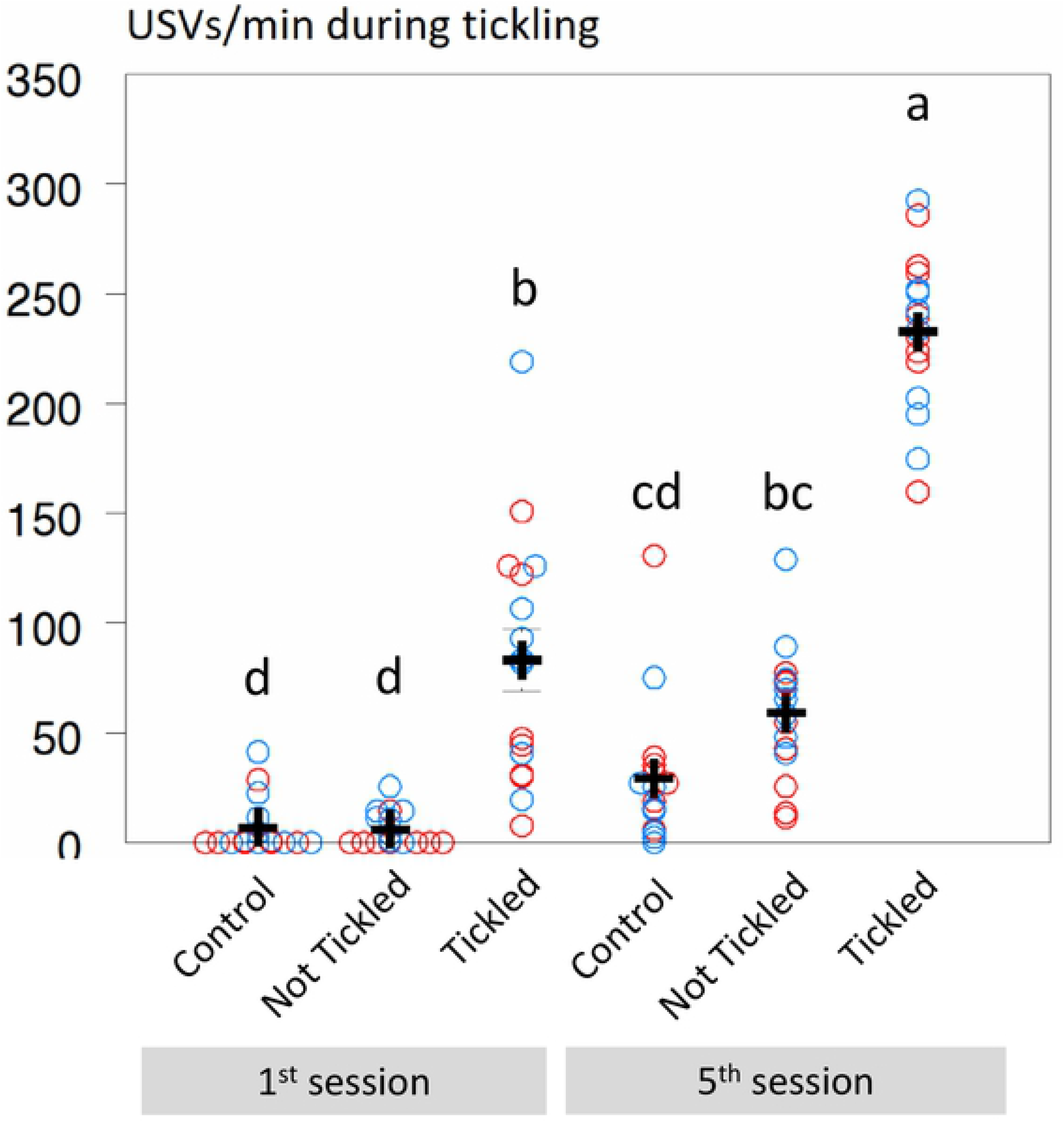
Data plot of 50 kHz ultrasonic vocalisations (USVs) per minute for individual rats during the 1^st^ and 5^th^ sessions in the periods when tickling occurred for the Tickled rats; the Control rats and the Not Tickled rats were not tickled during these periods (red circles: Odour A; blue circles: Odour B). Means (± s.e.) are indicated with black plus symbols for each grouping, and different superscripts indicate significant difference (P < 0.001).

**Figure 2b.**
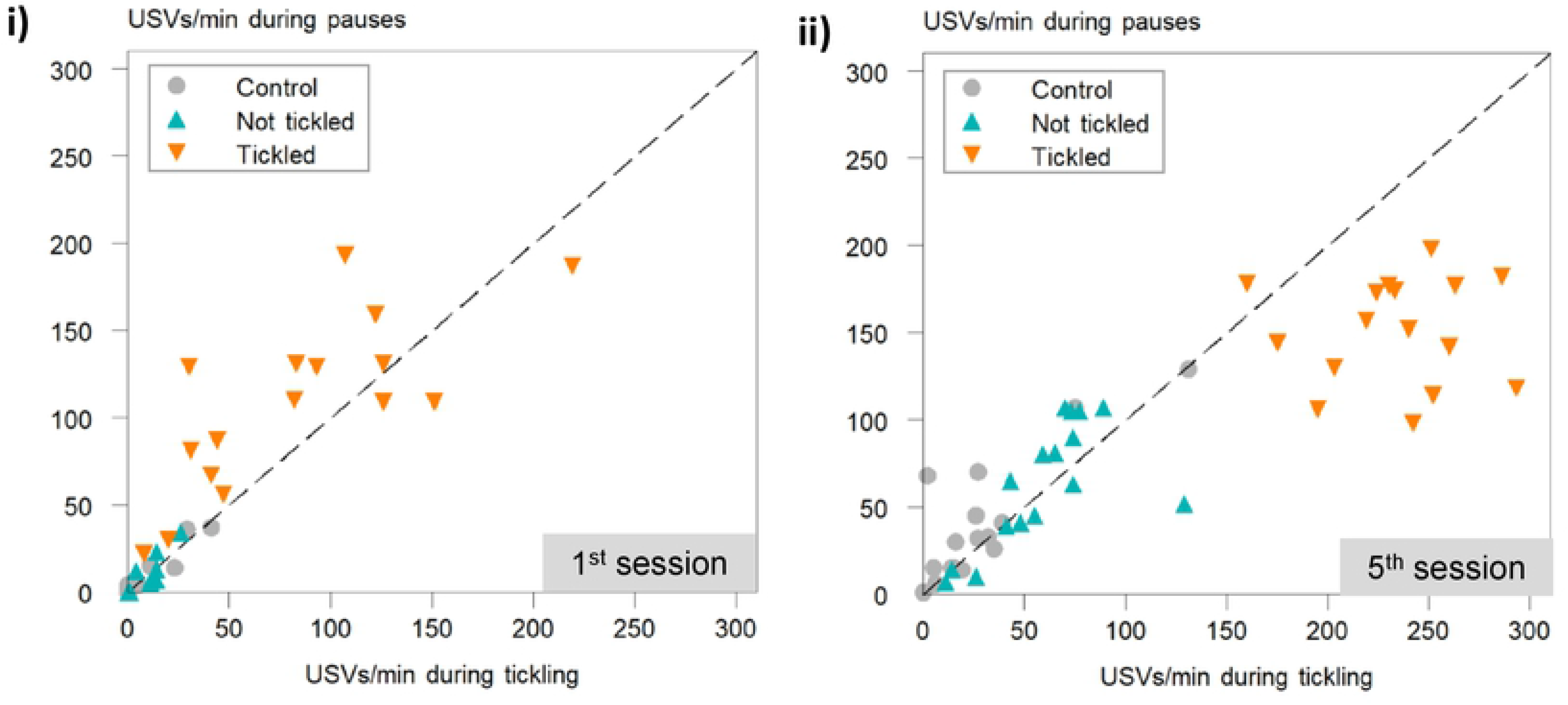
Data of 50 kHz ultrasonic vocalisations (USVs) per minute for individual rats during the pauses plotted against 50 kHz USVs per minute in the periods when tickling occurred for the Tickled rats for the **i)** 1^st^ and **ii)** 5^th^ sessions; the Control rats and the Not Tickled rats were not tickled during these periods. The dashed lines indicate where Y=X.

Most rats also emitted 50 kHz USVs during the pauses between tickling periods. When rats were tickled, although numerically greater, the USVs during pauses did not differ significantly from those observed during the tickling in the 1^st^ session (108 vs 83 (±11.1) USVs/min; P = 0.390), whereas by the 5^th^ session, significantly more 50 kHz USVs were emitted during tickling that during the pauses (151 vs 233 (±11.1) USVs/min for pauses and tickling periods, respectively; P < 0.001; Figure 2b). For the Not Tickled rats, including Controls, the levels of 50 kHz USVs were similar between tickling periods and pauses because no actual tickling took place. When rats were tickled, 50 kHz USVs emitted during pauses and during tickling were correlated for the 1^st^ (Pearson’s r=0.76; P = 0.001) but not the 5^th^ session (Pearson’s r=0.02; P = 0.955). Medians of the behaviour during pauses in the 5^th^ session are shown in Table 1, with tickled rats showing significantly more hand seeking behaviour and play jumping, with consequently less time spent in general locomotion, than when Not Tickled or compared to Control rats.

**Table 1.**
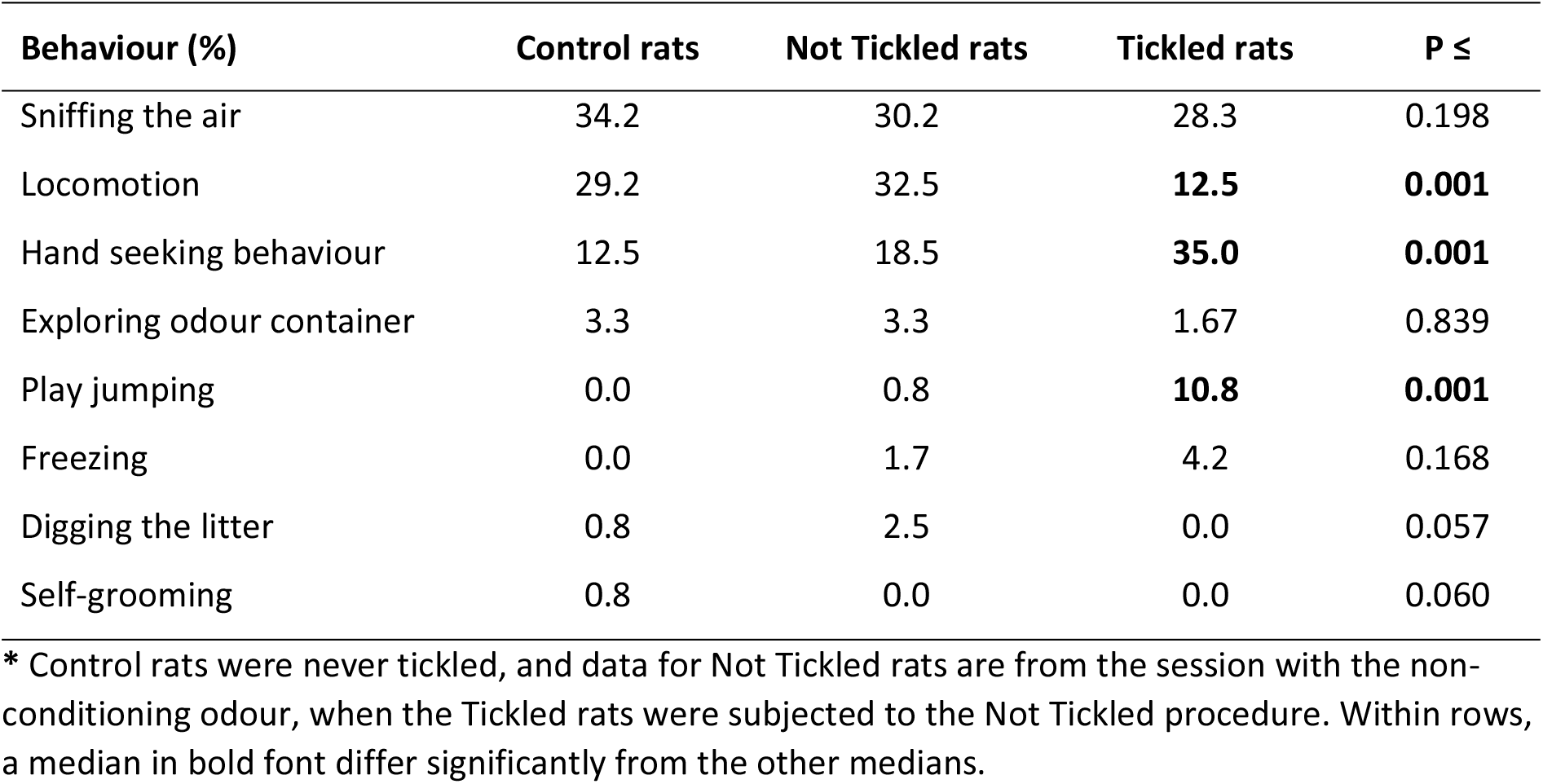
Median percentage of time spent in different behaviours by rats during pauses between tickling* periods in the 5^th^ conditioning session.

Even after five tickling sessions, rats did not emit more anticipatory USVs when exposed to their tickling odour, compared to when exposed to their non-tickling odour and compared to the control group (F_2,42_=0.33; P=0.718; Figure 3). Figure 4 shows the frequency of 50 kHz USVs emitted by the rats as a function of their anticipatory 50 kHz USVs. The higher level of USVs during tickling is clearly visible, but with no clear correlation with anticipatory USVs for the tickled rats. However, for the sessions without tickling (Not Tickled and Control rats), the 50 kHz USVs emitted show a positive relationship with the 50 kHz USVs emitted during the pre-session (anticipatory) minute (R^2^=53.1%; T=4.99: P <0.001; Figure 4), supporting the previous finding that rats may be characterised according to their level of vocalisation.

**Figure 3.**
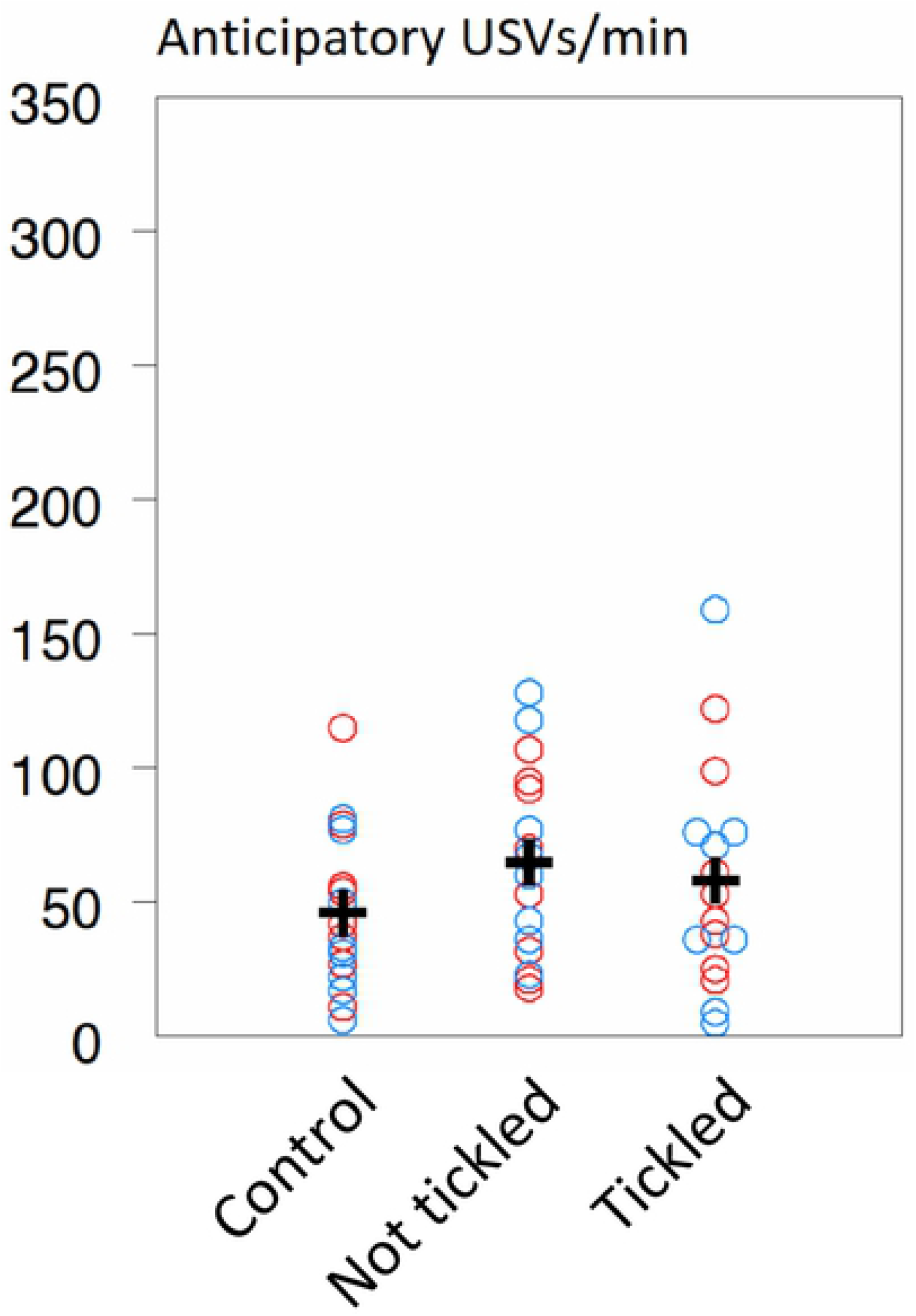
Data plot of 50 kHz ultrasonic vocalisations (USVs) per minute for individual rats during the 5^th^ session from the 60 s period (anticipatory) prior to tickling; the Control rats and the Non-tickled rats were not tickled during the sessions that followed (red circles: Odour A; blue circles: Odour B). Means (± s.e.) are indicated with black plus symbols for each grouping, and the y-axis scale is comparable to Figure 2a.

**Figure 4.**
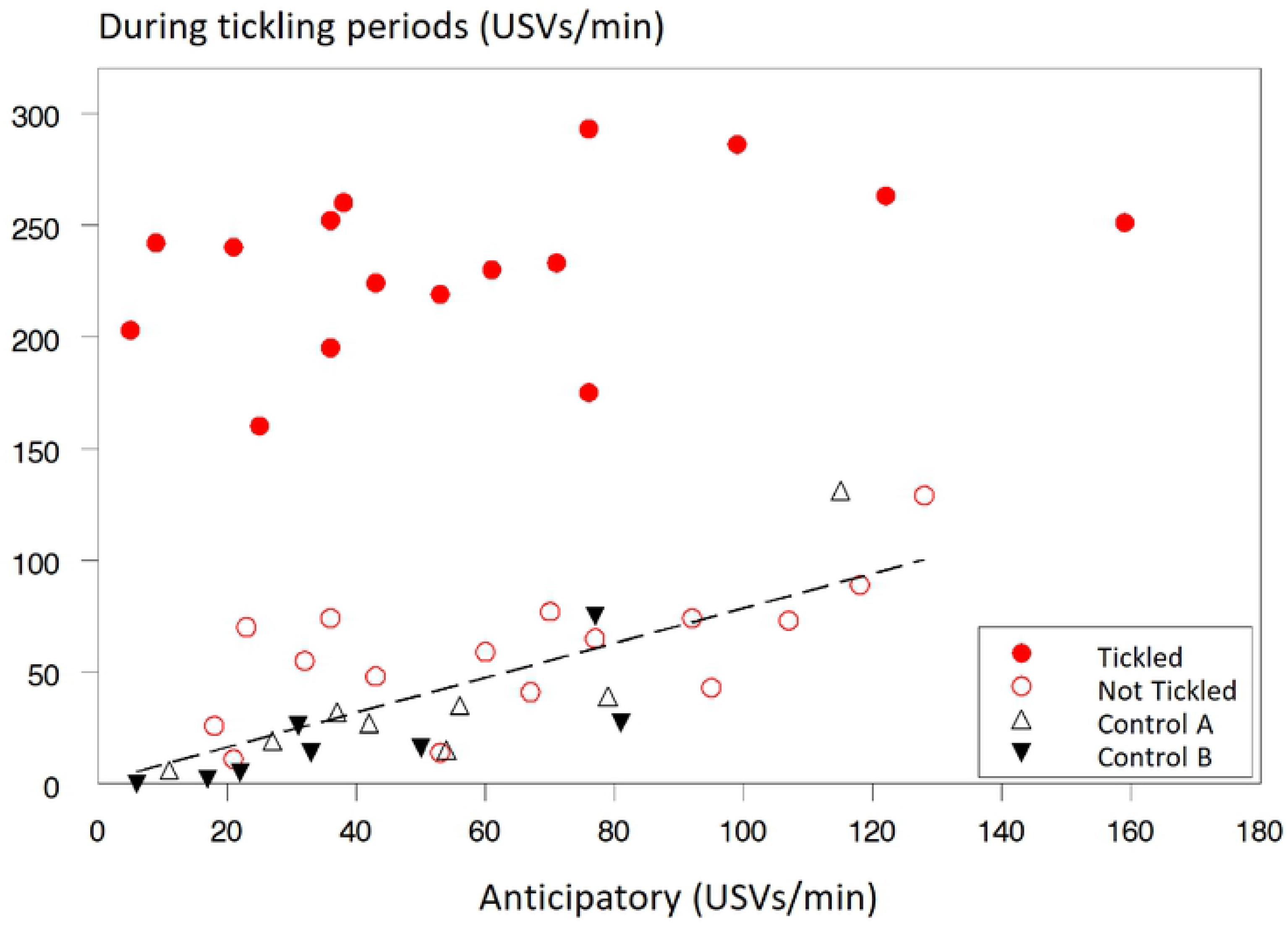
Data plot of 50 kHz ultrasonic vocalisations (USVs) per minute during the 5^th^ session in the periods when tickling occurred for the Tickled rats plotted against the USVs per minute during the 60-s period (anticipatory) prior to tickling. Each data point is an individual rat, with Control rats plotted for both Odours A and B. The regression equation, where data from Tickled rats have been excluded, is Y = 1 + 0.78X (R^2^=58%; P < 0.001).

USVs emitted during the Triple Odour test are shown in Figure 5. During the first minute, where no odour was present, the frequencies of 50 kHz USVs were no different from those seen during the anticipatory period during odour condition (see Figure 3). Over the course of the Triple odour test, USV frequency decreased gradually, but a significant increase in 50 kHz USVs from the preceding pause was found for the tickled rats when exposed to their tickling odour (increase: 19, 31 and −3 (±5.9) USVs/min for Tickled A, Tickled B, and Control rats, respectively; F_2,21_=8.7; P = 0.002), with the increase being significantly different for both A-tickled (P=0.032) and B-tickled rats (P=0.001) from that of the controls. The rats thus increase their USV frequency when their conditioning odour was presented indicating that the rats had learned to associate an odour with the positive experience of tickling.

**Figure 5.**
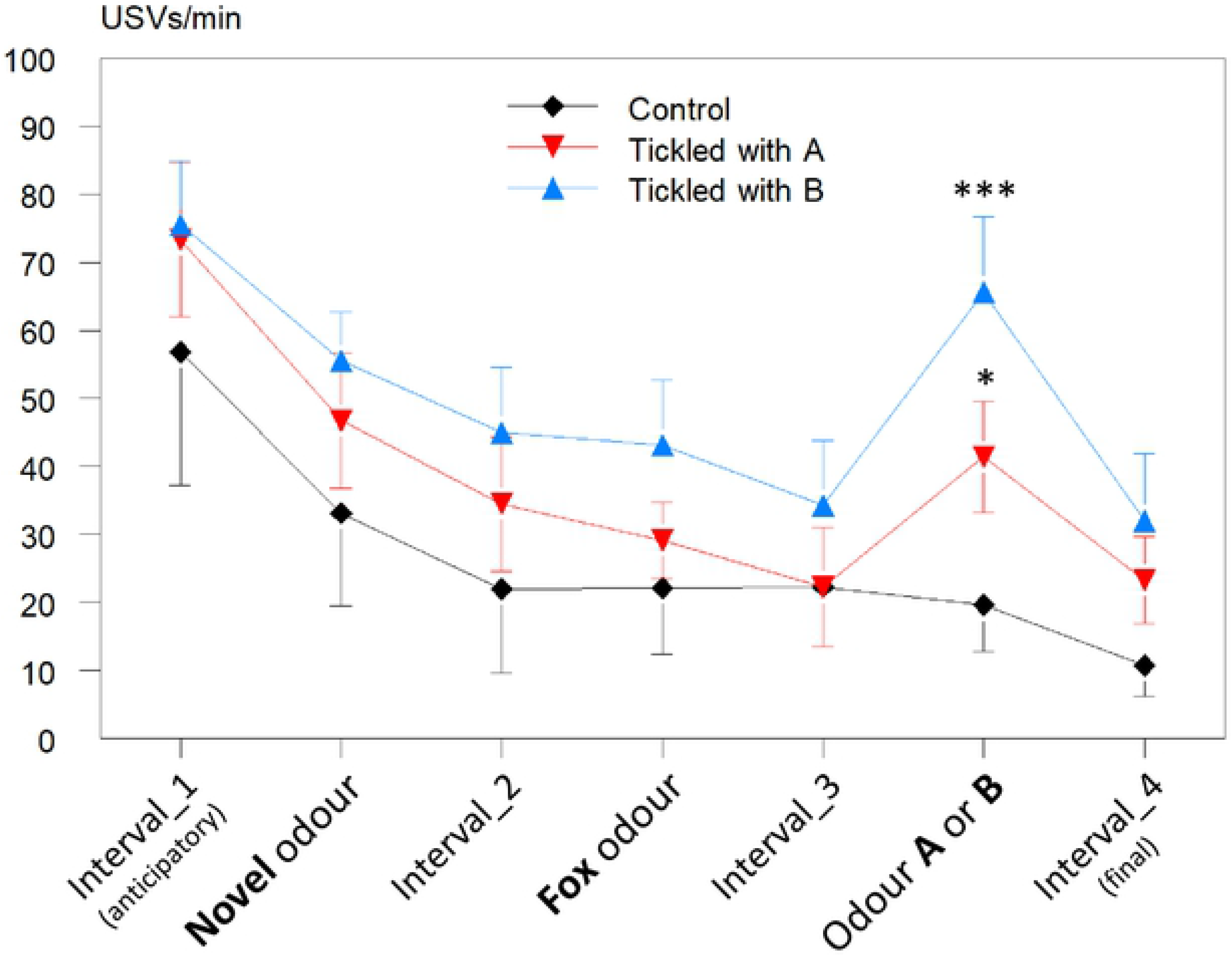
Mean number of 50 kHz ultrasonic vocalisations (USVs ± s.e.) per minute during the Triple Odour test for Tickled A, Tickled B, and Control rats. Tickled rats were exposed to their conditioning odour as the third odour, and for the Control rats, half were exposed to Odour A and half to Odour B. Asterisks indicate a significant increase in USVs (* P = 0.032; *** P = 0.001).

Freezing was scored during the Triple Odour test, as exposure to fox odour was expected to induce more freezing as an indicator of fear. However, as shown in Figure 6, freezing did not increase when fox odour was in the arena. The Tickled rats showed the same low level of freezing throughout the test, independent of odour present. The Control rats, however, showed a significant increase in freezing when one of the conditioned odours were present, and this was mainly due to greatly elevated levels of freezing in the Control rats (n=4) exposed to odour A (F_3,19_=9.0; P=0.001; Figure 6).

**Figure 6.**
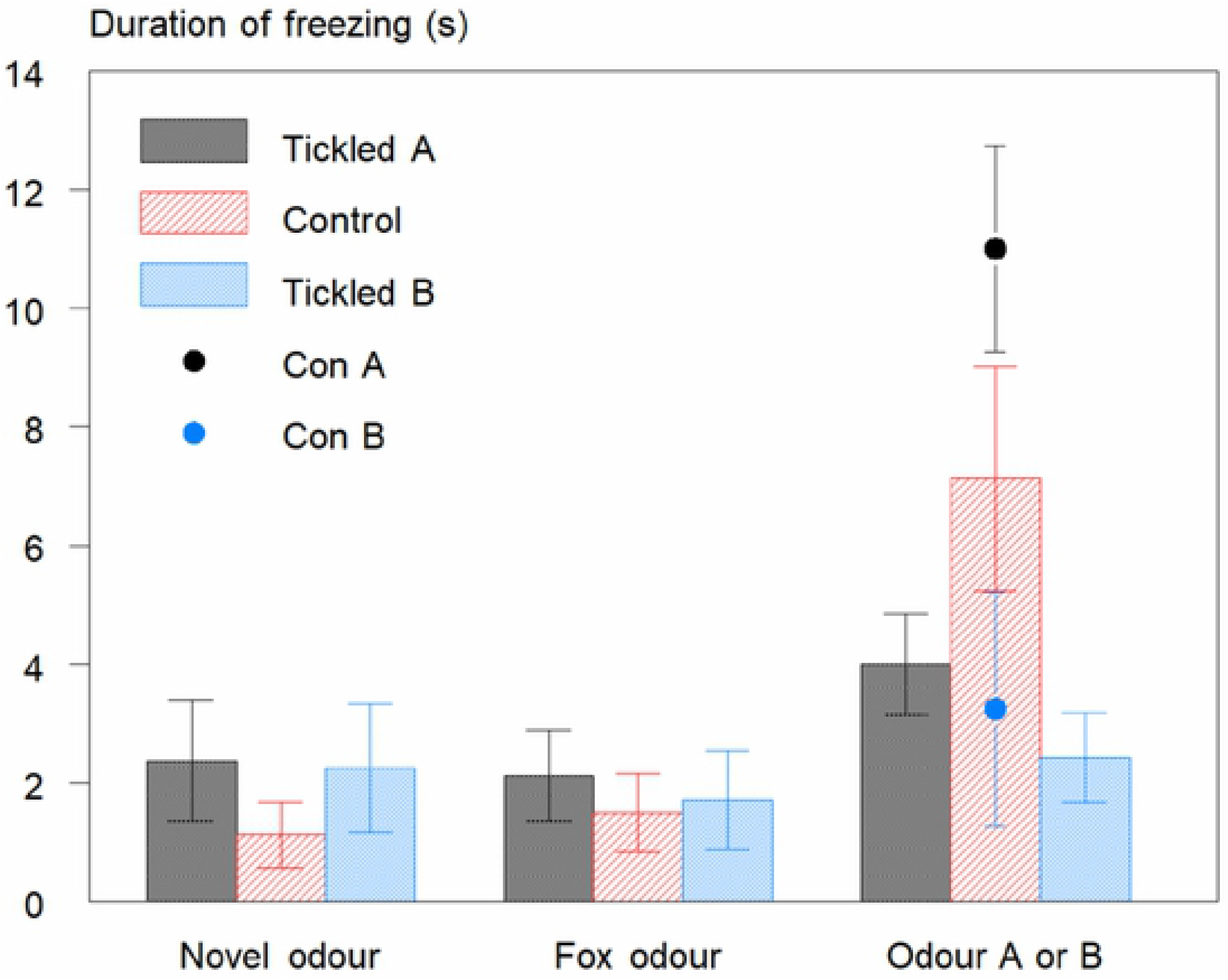
Mean duration (s ± s.e.) of freezing behaviour during exposure to the three odours in the Triple Odour test. Bars indicate the mean for each treatment group (n=8). For the third odour exposure (Odour A or B), the means for Control rats are also shown separately for rats exposed to Odour A (Con A; n=4) and to Odour B (Con B; n=4).

Behaviour during the T-maze test showed that overall, more time was spent in the arm with odour A (Figure 7). This was significantly different from time spent in the arm with odour B for the Tickled A rats (F1,14=5.0; P = 0.041) with a similar tendency for Control rats (P = 0.088). The proportion of time spent in the arm with Odour A was significantly different from chance (0.5) only for the rats tickled with odour A (0.60; T=2.51; P = 0.041).

**Figure 7.**
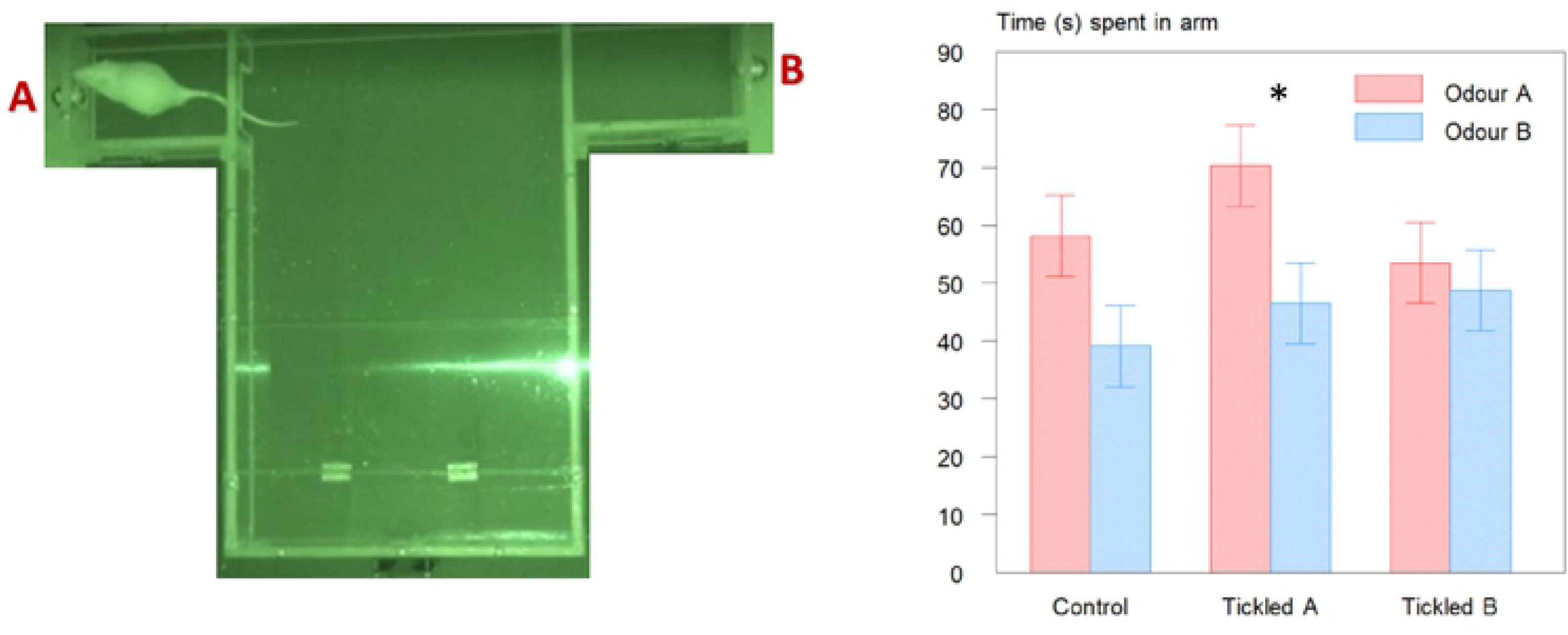
Time (s; mean ± s.e.) spent in each arm of the T-maze for Tickled and Control rats. Asterisk indicate a significant difference between arms within treatment group (* P = 0.041). Photo shows a screen-shots from the video recording of a test, with the rat in one of the two arms. Each arm was fitted with a tea-ball containing one of the two odours. The ventilator drawing air from both arms into the main area can be seen at the bottom of the photo.

## General discussion

Using an appetitive conditioning method, we aimed to condition rats to learn to associate the presence of an odour with the positive experience of tickling. Our first hypothesis was that rats, which had learned to make the odour-tickling association would emit more anticipatory USVs when exposed to the odour prior to being tickled. This was not the case, as no differences in anticipatory USVs were found between the treatment groups. Indeed, our use of the term anticipatory can be questioned, given the findings. Heyse et al. (2015) found that rats would emit 50 kHz calls in anticipation of access to a running wheel. Others have found that individual rats vocalize more in a chamber associated with play than in a habituated control chamber (Knutson et al., 1998), indicative of anticipation of positive experiences. The absence of similar anticipatory vocalisations in the present experiment would indicate that, with respect to our first hypothesis, the conditioning paradigm was not successful. However, the levels of USVs emitted during the (anticipatory) pre-session minute appeared to predict the overall level of vocalisation for individual rats when these were not tickled, suggesting that rats can be categorised according to their USV frequency independent of any tickling occurring. This is in accordance with Burgdorf et al. (2005, 2013), who divergently selected rats based on their 50 kHz vocalisations. It has previously been found that tickling-induced 50 kHz ultrasonic vocalizations are individually stable and can predict behaviour in tests of anxiety and depression in rats (Schwarting et al. 2007; Mällo et al., 2007). Our finding that USVs produced when not being tickled show large inter-individual differences, but little intra-individual variation, is complementary to these results.

The second hypothesis was that more USVs would be emitted by the tickled than by control rats when exposed to the conditioned odour following exposure to an aversive odour. We found an increase in USVs produced by the tickled rats when their tickling odour was placed in the arena.

Given that 50 kHz USVs are indicative of positive affect (e.g. Burgdorf and Panksepp, 2006), this would indicate that the tickled rats had learned to associate the odour with a positive experience. Ideally, we would have tested the conditioned rats with both odours, but the small number of animals made this statistically inappropriate. However, the increase in 50 kHz USVs by the Tickled A and Tickled B rats when exposed to their conditioned odour in the Triple Odour test was not simply because the odour was known compared to the two previous odours, as the Control rats showed no such increase in USV production. It was noted that exposure to the fox odour did not provoke neither 22 kHz vocalisations, nor freezing behaviour in the rats, indicating that this odour was less aversive than anticipated.

We also expected tickled rats to spend more time in the arm of a T-maze containing their tickling odour. This was found only for rats tickled in the presence of odour A. However, control rats appeared to be more attracted to odour A, and for the rats tickled while exposed to odour B, this preference for odour A was not evident. This could be an indication that they had developed an attraction to odour B, which was sufficiently strong to eliminate a potentially intrinsic preference for odour A. This corresponds to the findings from the Triple Odour test, where odour B appeared to have a stronger effect than odour A (see Figure 5). Control rats also showed more freezing when exposed to odour A in the Triple Odour test. Although freezing is often considered an indication of fear, the behaviour is but a display of increased alertness, and the interpretation is context specific. Exposure to oestrus odours can elicit freezing in rats (Nielsen et al., 2013, 2019) and this is enhanced by sexual experience (Nielsen et al., 2016). It may be that odour A induced freezing in control rats because they find it more interesting, as shown by their un-conditioned preference in the T-maze test. The two odours were chosen as being neutral to rats (Devore et al., 2013), and we struggle to explain why they affect the behaviour of the rats differently. One, speculative possibility is a potential sedative effect of inhaling limonene resulting in decreased locomotor activity, which has been found in mice (Carvalho-Freitas and Costa, 2002; Wolffenbüttel et al., 2018), but no differences in activity of the rats were found among treatments during odour conditioning.

As mentioned in the introduction, aversive conditioning is a widely used technique in learning studies of memory and other brain functions in laboratory rodents (e.g. Ellis and Kesner, 1983; Chapuis et al., 2007; Raineki et al., 2009). Pairing aversive stimuli such as electric shocks, with a neutral stimulus or situation usually give rise to associations learned within a few sessions (Kroon and Carobrez, 2009). In contrast, the application of appetitive conditioning regimes often require more pairing sessions to become effective. Although our positive conditioning was successful, as shown by the response of the rats to their conditioning odour in the Triple Odour test, this did not appear to be a very strong association, as no increase in anticipatory vocalisation was seen, nor a very convincing preference for the tickling odour in the T-maze test. Others have also struggled to demonstrate a link between increased 50 kHz USVs and reward-related stimuli: Brenes and Schwarting (2014) conditioned rats to associate a tone with a food reward, and measured the expression of reward anticipation as increases in USVs. However, when the rats were food-deprived, they showed only behavioural but not vocal anticipation and when sated, the reward cue continued to elicit 50 kHz USVs despite being devalued by pre-feeding. These findings may, in part, have been due to the large inter-individual variability among rats, giving rise to different types of responders (Brenes and Schwarting, 2015). It is evident from these and the present results that appetitive conditioning is likely to be more complex and less effective than most aversive conditioning.

One protocol of rat tickling has been described in detail by Cloutier et al. (2018). The benefit of this is that it allows comparisons to be made if the same method is used across experiments. However, we did not standardise the tickling method used in the present experiment, over and above the fixed alternating periods of tickling and pauses. This was a conscious choice on our part, as we had previously found a large individual variation in the response of the rats to tickling. Our experience indicated that this variation was reduced if the rats were tickled and played with whilst allowing the hand to react to the behavioural responses of the individual rat. In addition, as tickling is a playful experience, it should be varied and unpredictable to the rats. Although the lack of standardisation prevented us from comparing behaviour of the rats during the active tickling period, i.e. as the behaviour of the experimenter varied slightly across rats and across sessions, we were able to use the hand-seeking behaviour during the pauses to assess the likability of tickling for each rat. Tickled rats showed more hand seeking behaviour and play jumping with simultaneously more 50 kHz USVs emitted during the pauses between tickling, indicating that the tickling lead to a positive affective state.

In conclusion, rats learned to associate an odour with the positive experience of being tickled, as they increased their 50 kHz USVs when exposed to this odour in a test situation without tickling, compared to control rats that had been exposed to the same odour for the same amount of time without being tickled. However, no increase was seen in anticipatory USVs when exposed to the conditioning odour prior to being tickled, and only one of the conditioning odours gave rise to a preference by the tickled rats in a T-maze test. These findings indicate that rats can learn to associate an odour with the positive experience of tickling, and positive odour conditioning may thus have potential to be developed further with a view to replacing negative odour conditioning tests. However, different odours may differ in their efficacy, and appetitive (positive) conditioning is clearly more difficult and slower to induce than aversive (negative) conditioning.

## Acknowledgements

The authors are grateful to the staff at the animal facility for taking care of the rats. This study was supported by the RESAS Strategic Research Programme and the Roslin Institute (University of Edinburgh) BBSRC Institute Strategic Programme, as well as a Credits Incitatifs grant from the Department of Animal Physiology and Livestock Systems (PHASE), INRA, France.

## References

Avvisati R, Contu L, Stendardo E, Michetti C, Montanari C, Scattoni ML, Badiani A. Ultrasonic vocalization in rats self-administering heroin and cocaine in different settings: evidence of substance-specific interactions between drug and setting. Psychopharmacol. 2016;233: 1501–1511. doi: 10.1007/s00213-016-4247-4.

Barker DJ. (2018). Ultrasonic vocalizations as an index of positive emotional state. In: Brudzynski SM, editor. Handbook of Behavioral Neuroscience. 2018. pp. 253–260. doi: 10.1016/b978-0-12-809600-0.00024-x.

Blanchard RJ, Blanchard DC, Agullana R, Weiss SM. Twenty-two kHz alarm cries to presentation of a predator, by laboratory rats living in visible burrow systems. Physiol Behav. 1991;50: 967–972. doi: 10.1016/0031-9384(91)90423-L.

Boissy A, Manteuffel G, Jensen MB, Moe RO, Spruijt B, Keeling LJ, Winckler C, Forkman B, Dimitrov I, Langbein J, Bakken M, Veissier I, Aubert A. Assessment of positive emotions in animals to improve their welfare. Physiol Behav. 2007;92: 375–397. doi: 10.1016/j.physbeh.2007.02.003.

Brenes JC, Schwarting RKW. Attribution and expression of incentive salience are differentially signaled by ultrasonic vocalizations in rats. PLoS ONE. 2014;9: e102414. doi: 10.1371/journal.pone.0102414.

Brenes JC, Schwarting RKW. Individual differences in anticipatory activity to food rewards predict cue-induced appetitive 50-kHz calls in rats. Physiol. Behav. 2015;149: 107–118. doi: 10.1016/j.physbeh.2015.05.012.

Burgdorf J, Panksepp J. The neurobiology of positive emotions. Neurosci Biobehav Rev. 2006;30: 173–187. doi: 10.1016/j.neubiorev.2005.06.001.

Burgdorf J, Moskal JR, Brudzynski SM, Panksepp J. Rats selectively bred for low levels of play-induced 50kHz vocalizations as a model for Autism Spectrum Disorders: A role for NMDA receptors. Behav. Brain Res. 2013;251: 18–24. doi: 10.1016/j.bbr.2013.04.022.

Burgdorf J, Panksepp J, Moskal JR. Frequency-modulated 50kHz ultrasonic vocalizations: A tool for uncovering the molecular substrates of positive affect. Neurosci Biobehav Rev. 2011;35: 1831–1836. doi: 10.1016/j.neubiorev.2010.11.011.

Burgdorf J, Panksepp J, Moskal JR. Rat 22-kHz ultrasonic vocalizations as a measure of emotional set point during social interactions. In: Brudzynski SM, editor. Handbook of Behavioral Neuroscience. 2018. pp. 261–265. doi: 10.1016/b978-0-12-809600-0.00025-1.

Burgdorf J, Panksepp J, Brudzynski SM, Kroes R, Moskal JR. Breeding for 50-kHz positive affective vocalization in rats. Behav Genet. 2005;35: 67–72. doi:10.1007/s10519-004-0856-5.

Caffrey MK, Febo M. Cocaine-associated odor cue re-exposure increases blood oxygenation level dependent signal in memory and reward regions of the maternal rat brain. Drug Alcohol Depend. 2014;134: 167–177. doi: 10.1016/j.drugalcdep.2013.09.032.

Carvalho-Freitas MIR, Costa M. Anxiolytic and sedative effects of extracts and essential oil from *Citrus aurantium L*. Biol Pharmaceut Bull. 2002;25: 1629–1633. doi: 10.1248/bpb.25.1629.

Chapuis J, Messaoudi B, Ferreira G, Ravel N. Importance of retronasal and orthonasal olfaction for odor aversion memory in rats. Behav Neurosci. 2007;121: 1383–1392. DOI: 10.1037/0735-7044.121.6.1383.

Choi J-S, Brown TH. Central amygdala lesions block ultrasonic vocalization and freezing as conditional but not unconditional responses. J Neurosci. 2003;23: 8713–8721. PubMed ID: 14507971.

Cloutier S, LaFollette MR, Gaskill BN, Panksepp J, Newberry RC. Tickling, a technique for inducing positive affect when handling rats. J Vis Exp. 2018;135: e57190. doi: 10.3791/57190.

Deehan GA Jr, Ding Z-M, Hauser SR, Engleman EA, Wilden JA, McBride WJ, Rodd ZA. Conditioned odor cues associated with the access to or the absence of alcohol differentially modulate dopmamine efflux in the nucleus accumbens shell. Alcohol Clin Exp Res. 2012; 36: 186A.

Devore S, Lee J, Linster C. Odor preferences shape discrimination learning in rats. Behav Neurosci. 2013;127: 498–504. doi: 10.1037/a0033329.

Ellis ME, Kesner RP. The noradrenergic system of the amygdala and aversive information processing. Behav Neurosci. 1983;97: 399–415. doi: 10.1037//0735-7044.97.3.399.

Heyse NC, Brenes JC, Schwarting RK. Exercise reward induces appetitive 50-kHz calls in rats. Physiol Behav. 2015;147: 131–40. doi: 10.1016/j.physbeh.2015.04.021.

Knutson B, Burgdorf J, Panksepp J. Ultrasonic vocalizations as indices of affective states in rats. Psychol Bull. 2002;128: 961–977. doi: 10.1037//0033-2909.128.6.961.

Knutson B, Burgdorf J, Panksepp J. Anticipation of play elicits high-frequency ultrasonic vocalizations in young rats. J Comp Psychol. 1998;112: 65. doi: 10.1037/0735-7036.112.1.65.

Kroon JAV, Carobrez AP. Olfactory fear conditioning paradigm in rats: Effects of midazolam, propranolol or scopolamine. Neurobiol Learn Mem. 2009;91: 32–40. doi: 10.1016/j.nlm.2008.10.007.

LaFollette MR, O’Haire ME, Cloutier S, Blankenberger WB, Gaskill BN. Rat tickling: A systematic review of applications, outcomes, and moderators. PLoS ONE. 2017;6: 12:e0175320. doi: 10.1371/journal.pone.0175320.

LaFollette MR, O’Haire ME, Cloutier S, Gaskill BN. Practical rat tickling: Determining an efficient and effective dosage of heterospecific play. Appl Anim Behav Sci. 2018;208: 82–91. doi: 10.1016/j.applanim.2018.08.005.

Lam H. Can odour be associated with positive affective states in rats (Rattus norvegicus)? MSc thesis, University of Edinburgh; 2017. 45 pp.

Lawrence AB, Newberry RC, Špinka M. Positive welfare: What does it add to the debate over pig welfare? In: Špinka M., editor. Advances in Pig Welfare. 2017. pp. 415–444. doi: 10.1016/b978-0-08-101012-9.00014-9.

Lowen SB, Rohan ML, Gillis TE, Thompson BS, Wellons CB, Andersen SL. Cocaine-conditioned odor cues without chronic exposure: Implications for the development of addiction vulnerability. Neuroimage Clin. 2015;8: 652–659. doi: 10.1016/j.nicl.2015.06.012.

Mällo T, Matrov D, Herm L, Kõiv K, Eller M, Rinken A, Harro J. Tickling-induced 50-kHz ultrasonic vocalization is individually stable and predicts behaviour in tests of anxiety and depression in rats. Behav Brain Res. 2007;184: 57–71. doi: 10.1016/j.bbr.2007.06.01.

Moriceau S, Wilson DA, Levine S, Sullivan RM. Dual circuitry for odor-shock conditioning during infancy: Corticosterone switches between fear and attraction via amygdala. J Neurosci. 2006;26: 6737–6748. doi: 10.1523/jneurosci.0499-06.2006.

Nielsen BL, Jerôme N, Saint-Albin A, Rampin O, Maurin Y. Behavioural response of sexually naïve and experienced male rats to the smell of 6-methyl-5-hepten-2-one and female rat faeces, Physiol Behav. 2013;120: 150–155. doi: 10.1016/j.physbeh.2013.07.012.

Nielsen BL, Jerôme N, Saint-Albin A, Ouali C, Rochut S, Zins E-L, Briant C, Guettier E, Reigner F, Couty I, Magistrini M, Rampin O. Oestrus odours from rats and mares: behavioural responses of sexually naive and experienced rats to natural odours and odorants. Appl Anim Behav Sci. 2016;176: 128–135. doi: 10.1016/j.applanim.2016.01.014 10.1016/0003-3472(86)90014-x.

Nielsen BL, Jérôme N, Saint-Albin A, Joly F, Rabot S, Meunier N. Sexual responses of male rats to odours from female rats in oestrus are not affected by female germ-free status. Behav Brain Res. 2019; 359: 686–693. doi: 10.1016/j.bbr.2018.09.018.

Panksepp J, Burgdorf J. Laughing rats? Playful tickling arouses high-frequency ultrasonic chirping in young rodents. Am J Play. 2010;2: 357–372.

Popik P, Kos T, Pluta H, Nikiforuk A, Rojek K, Rygula R. Inhibition of the glucocorticoid synthesis reverses stress-induced decrease in rat’s 50-kHz ultrasonic vocalizations. Behav Brain Res. 2014;260: 53–57. doi: 10.1016/j.bbr.2013.11.029.

Raineki C, Shionoya K, Sander K, Sullivan RM. Ontogeny of odor-LiCl vs. odor-shock learning: Similar behaviors but divergent ages of functional amygdala emergence. Learn Mem. 2009;16: 114–121. doi: 10.1101/lm.977909.

Revillo DA, Fernandez G, Castello S, Paglini MG, Arias C. Odor-avoidance or odor-preference induced by amphetamine in the infant rat depending on the dose and testing modality. Behav Brain Res. 2012;231: 201–207. doi: 10.1016/j.bbr.2012.03.018.

Rygula R, Pluta H, Popik P. Laughing rats are optimistic. PLoS ONE 2012;7: e51959. doi: 10.1371/journal.pone.0051959.

Schwarting RK, Jegan N, Wöhr M. Situational factors, conditions and individual variables which can determine ultrasonic vocalizations in male adult Wistar rats. Behav Brain Res. 2007;182: 208–222. doi: 10.1016/j.bbr.2007.01.029.

Shide DJ, Blass EM. Opioid mediation of odor preferences induced by sugar and fat in 6-day-old rats. Physiol Behav. 1991;50: 961–966. doi: 10.1016/0031-9384(91)90422-k.

Sullivan RM, Wilson DA, Ravel N, Mouly AM. Olfactory memory networks: from emotional learning to social behaviors. Front Behav Neurosci. 2015;9: 36. doi: 10.3389/fnbeh.2015.00036.

Tonoue T, Ashida Y, Makino H, Hata H. Inhibition of shock-elicited ultrasonic vocalization by opioid peptides in the rat: a psychotropic effect. Psychoneuroendocrinol. 1986; 11: 177–184. doi: 10.1016/0306-4530(86)90052-1.

Torquet N, Aime P, Messaoudi B, Garcia S, Ey E, Gervais R, Julliard AK, Ravel N. Olfactory preference conditioning changes the reward value of reinforced and non-reinforced odors. Front Behav Neurosci. 2014;8: 29. doi: 10.3389/fnbeh.2014.00229.

Wöhr M, Schwarting RK. Affective communication in rodents: ultrasonic vocalizations as a tool for research on emotion and motivation. Cell Tissue Research 2013;354: 81–97. doi: 10.1007/s00441-013-1607-9

Wolffenbüttel AN, Zamboni A, Becker G, dos Santos MK, Borille BT, de Cássia Mariotti K, Fagundes AC, de Oliveira Salomón JL, Coelho VR, Ruiz LV, de Moura Linck V, Dallegrave E, Cano P, Esquifino AI, Leal MB, Limberg RP. Citrus essential oils inhalation by mice: Behavioral testing, GCMS plasma analysis, corticosterone, and melatonin levels evaluation. Phytother Res. 2018;32: 160–169. doi: 10.1002/ptr.5964.

